# Multiscale simulation of peripheral neural signaling

**DOI:** 10.1101/196451

**Authors:** CH Lubba, Y Le Guen, S Jarvis, NS Jones, SC Cork, A Eftekhar, SR Schultz

**Affiliations:** Centre for Neurotechnology, Imperial College London, South Kensington, London SW7 2AZ, UK; Department of Bioengineering, Imperial College London, South Kensington, London SW7 2AZ, UK; Department of Mathematics, Imperial College London, South Kensington, London SW7 2AZ, UK; Centre for Mathematics of Precision Healthcare, Imperial College London, South Kensington, London SW7 2AZ, UK; Department of Medicine, Imperial College London, South Kensington, London SW7 2AZ, UK; Department of Electrical and Electronic Engineering, Imperial College London, South Kensington, London SW7 2AZ, UK

## Abstract

Bioelectronic Medicines that modulate the activity patterns on peripheral nerves have promise as a new way of treating diverse medical conditions from epilepsy to rheumatism. Progress in the field builds upon time consuming and expensive experiments in living organisms to evaluate spontaneous activity patterns, stimulation efficiency, and organ responses. To reduce experimentation load and allow for a faster, more detailed analysis of both recording from and stimulation of peripheral nerves, adaptable computational models incorporating insights won in experiments will be of great help. We present a peripheral nerve simulator that combines biophysical axon models and numerically solved and idealized extracellular space models in one environment. Two different scales of abstraction were merged. On the one hand we modeled the extracellular space in a finite element solver as a three dimensional resistive continuum governed by the electro-quasistatic approximation of the Maxwell equations. Potential distributions were precomputed for different media (homogeneous, nerve in saline, nerve in cuff). Axons, on the other hand, were modeled at a higher level of abstraction as one dimensional chains of compartments; each consisting of lumped linear elements and, for some, channels with non-linear dynamics. Unmyelinated fibres were based on the Hodgkin-Huxley model; for myelinated fibers, we instead adapted the model proposed by McIntyre et al. in 2002 to smaller diameters. To obtain realistic axon shapes, an iterative algorithm positioned fibers along the nerve with variable tortuosity, with tortuosity values fit to imaged trajectories. We validated our model with data from the stimulated rat vagus nerve. Simulation results predicted that tortuosity leads to differentiation in recorded signal shapes, with unmyelinated axons being the most affected. Tortuosity was further shown to increase the stimulation threshold. The model we developed can easily be adapted to different nerves, and may be of use for Bioelectronic Medicine research in the future.

## 1. Introduction

Manipulations of the peripheral nervous system (PNS) by implanted devices might soon serve as a treatment for various medical conditions. Such Bioelectronic Medicines[1] can be seen as a permanent, highly localised alternative to molecular medicines with less side effects. Already today, vagus nerve stimulation is being applied in patients suffering from refractory epilepsy [2], Alzheimer’s disease [3], anxiety[4], obesity [5], chronic heart failure [6], and to evoke anti-inflammatory effects [7, 8]. The more localized targeting of organs e.g. of the heart [9] has also shown promising results in animal experiments.

Current devices operate in open-loop mode and stimulation selectivity is low. Future Bioelectronic Medicines will need more precise stimulation interfaces and the capability to analyze (or ‘decode’) nerve activity to stimulate in an adaptive manner. First advances towards a decoding of information from peripheral nerves have been successfully undertaken [10, 11]. To both accelerate the design of interfaces and to further develop decoding algorithms, computational peripheral nerve models that integrate physiological insights won in experiments at different levels (i.e. properties of axons, extracellular media, spontaneous activity patterns, organ responses) will be of great merit – to predict stimulation efficiency and recording selectivity and as a source of surrogate data.

Previous efforts to simulate peripheral nerves date back approximately twenty years. In 1997, Struijk [12] developed a 2D model of recordings from myelinated peripheral axons. Models for stimulation were also proposed at the time [13, 14]. One main difficulty in peripheral nerve simulations, already appreciated at that time, is the calculation of extracellular potential from membrane currents in the inhomogeneous medium surrounding the axons. Early simulations therefore often concentrated on modeling the extracellular space while approximating axons with simplified models such as the Fitz-Hugh-Nagumo equations [e.g. 15] or the McNeal model [13, 16]. Only recently have precise biophysical axon models and detailed, numerically solved models of the surrounding medium been combined [17, 18], thanks to increasing availability of computational resources. Still, current models are limited in usability, adaptability and precision. They usually reside in two simulation environments: A compartmental simulator for axon models and a finite element methods (FEM) solver for the extracellular space. Both need to be coordinated in terms of geometry, coordinate systems, units, etc and setting up a model this way is a time consuming and error prone process that generally limits adaptability and the possibility of reusing results. Finally, fibres are usually modeled as being perfectly straight, which is an over simplification as we will show in the following.

In order to create a reusable, easily adaptable yet detailed simulation infrastructure where biophysical axon models and numerically solved extracellular space models meet along well defined interfaces, we developed a novel Python toolbox, `PNPy`‡. `PNPy` makes good use of existing simulation tools for axons and media and integrates their respective results in a three-dimensional model with a clearly structured information flow. Where possible, geometrical symmetries and separation of time and space were used to minimize the number of simulation steps needed. It uses the NEURON simulator [19] to model axon membrane processes at the scale of ion channels. Standard models for both myelinated and unmyelinated axons in the diameter range found in the periphery were implemented. Their trajectory was generated automatically with a variable degree of tortuosity. Inhomogeneities of the extracellular medium were taken into account via import of numerically precomputed potential fields in the FEM solver COMSOL Multiphysics 4.3 and idealized transmission functions fit to FEM results. The user can stay in Python to stimulate nerves and record from them *in silico*.

## 2. Methodology

### 2.1. Nerve Stimulation Experiments

Experiments were carried out in accordance with the Animals (Scientific Procedures) Act 1986 (United Kingdom) and Home Office (United Kingdom) approved project and personal licenses, and experiments were approved by the Imperial College Animal Welfare Ethical Review Board under project license PPL 70/7365. Male Wistar rats (body weight 350-400 g) were initially anaesthetised with isoflurane. Urethane was then slowly administered through a tail vein (20 mg kg^*-*^1). The left cervical vagus nerve was exposed and contacted with a stainless steel pseudo-tripolar hook electrode of pole distance 1 - 2 mm for stimulation. To record from the nerve, a bipolar platinum hook electrode (pole distance 2 - 3 mm) was then wrapped around the anterior branch of the subdiaphragmatic vagus nerve with an Ag/AgCl ground electrode placed in the abdominal cavity. Distance between both electrodes was 7 - 8 cm. Mineral oil was applied to each site to insulate the electrodes from environmental and proximal noise sources. Stimulation of the cervical vagus nerve was achieved using a Keithley 6221 current source, controlled by Standard Commands for Programmable Instruments (SCPI) via a custom built Matlab interface. The Keithley 6221 allows for a maximum current of 100 mA with a voltage compliance of 100 V for a limited pulse width of 0.1 ms. Bipolar cuff recordings were achieved with an Intan Technology RHD2000 system, using a 16-channel bipolar ended amplifier (RHD221). The obtained recordings were averaged over 10 runs and slightly wavelet-denoised (PyWavelet, 12 levels, soft threshold 1) with the Sym7 Wavelet as proposed by Diedrich et al. [20].

### 2.2. Imaging of Peripheral Nerve Tortuosity

To reproduce the morphology of axons, we imaged the vagus and sciatic nerves in mice using two photon fluorescence imaging. In the experiment, ChAT-Cre FLEX-VSFP 2.3 mice were euthanised by i.p. overdose of pentobarbital (150 mg kg^*-*^1). The pre-thoracic left and right vagus nerves were surgically exposed and 0.5 cm sections were removed and placed in phosphate buffered saline (155.1 mmol NaCl,2.96 mmol Na2HPO4, 1.05 mmol KH2PO4) adjusted to 8.0 pH with 1 mol NaOH. Sections of the left and right sciatic nerves of between 1 and 2 cm from above the knee were also removed. To prepare for microscopy, the nerves were placed on microscope slides, stretched until straight, and the nerve ends fixed with super glue. The preparation was covered with PBS. Distortions potentially caused by the stretching of the nerves were assumed to lie within the physiological range of movement-induced deformations the nerve undergoes in the living organism. A commercial 2P microscope was used for imaging (Scientifca, emission blue channel: 475/50 nm, yellow channel 545/55 nm, 511 nm dichroic, Semrock) while exciting at 950 nm using a Ti-Sapphire laser (Mai Tai HP, Spectra-Physics). All procedures were carried out in accordance with the Animals (Scientific Procedures) Act 1986 (United Kingdom) and Home Office (United Kingdom) approved project and personal licenses, and experiments were approved by the Imperial College Animal Welfare Ethical Review Board under project license PPL 70/7355.

### 2.3. PNPy Overview

Every `PNPy` simulation describes one peripheral nerve consisting of an arbitrary number of unmyelinated and myelinated axons, each with a certain diameter and trajectory. Axons can be activated by synaptic input, intra- and/ or extracellular stimulation. For extracellular recordings, electrodes are positioned along the nerve.

The module is organised as several core classes mapped to the physiological entities found in a peripheral nerve; shown in figure 2 along with the data flow. All objects are managed by the main class `Bundle`. This is the central object in the Python domain and represents the whole nerve. It contains instances of the `Axon`-class that defines properties needed by the NEURON simulations. Axon has two subclasses `Unmyelinated` and `Myelinated` for the two different types of fibres; each axon being characterised by its diameter and trajectory. To activate axons, `ExcitationMechanisms` are added to the `Bundle`. They can be either synaptic input (`UpstreamSpiking`), intracellular stimulation (`StimIntra`) or extracellular stimulation (`StimField`). Similarly for recording, electrodes can be added to the whole nerve as a `RecordingMechanism`. For all interactions with the extra-cellular space, i.e. extracellular stimulation or recording, a model of the medium defined in a `Extracellular`-class has to be set. This can be either homogeneous (`homogeneous`), an FEM result (`precomputedFEM`) or an analytically defined potential distribution (`analytic`).

**Figure 1:**
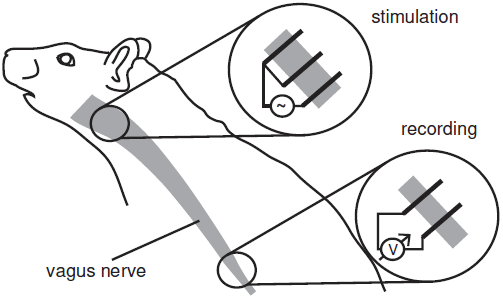
The validation data was obtained from a stimulated rat vagus nerve. A pseudotripolar electrode excited axons at the cervical vagus nerve, signals were picked up at the subdiaphragmatic vagus nerve with a bipolar electrode.

**Figure 2:**
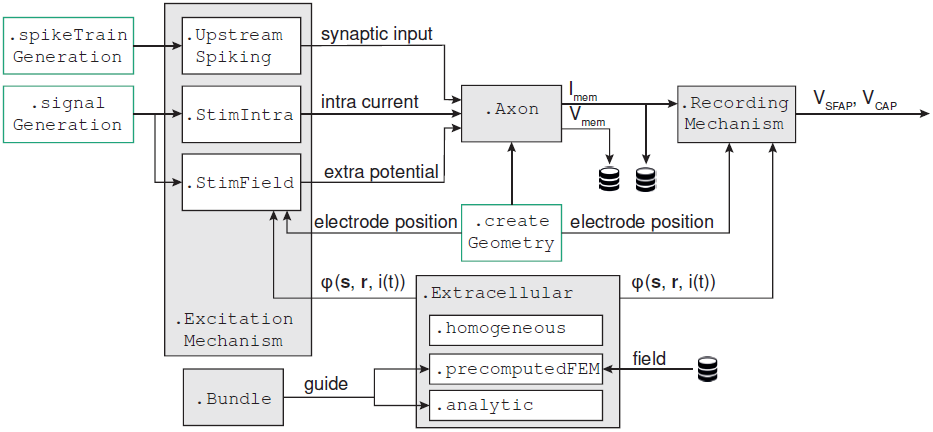
The `Axon`-class is the central object of `PNPys` internal information flow. Together with its associated `ExcitationMechanisms` it defines the NEURON simulation. `Extracellular`-objects allow the calculation of extracellular potentials given current *i*(*t*), source position s and receiver position **r**. They are used by both `StimField` for extracellular stimulation and by `RecordingMechanism` for recording. All classes are managed in the `Bundle`-class and supported by helper modules `spikeTrainGeneration`, `signalGeneration` and `createGeometry`.

In the simulation step, the definition of each axon in Bundle is sequentially transmitted to NEURON via the Python-NEURON-Interface [21] alongside its associated `ExcitationMechanisms`. After the calculation of single axon membrane processes is finished in NEURON, `PNPy` computes the extracellular single fiber action potential (SFAP) for the associated `RecordingMechanisms` from membrane currents. Once all axons have been processed, their contributions to the overall compound action potential (CAP) are added.

### 2.4. Assumptions and Simplifications

Several assumptions were required for the computational feasibility and efficiency of our model. Axons were assumed to be independent from each other in their activity. Properties such as diameter, myelination, and channel densities stayed constant along their length. The electro-quasistatic approximation of Maxwell’s equations governed the extracellular space, neglecting magnetic induction:

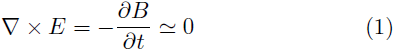

Further, all media were assumed to be purely resistive, so that all changes in current affected the potentials of the entire space immediately. In Maxwell’s equations this results in neglecting displacement currents:

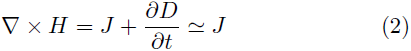

For the brain and in the considered frequency range, the electro-quasistatic approximation is assumed to be valid [22, 23]; previous peripheral nerve simulation studies have built on both approximations [12–14, 24]. Each tissue surrounding the nerve took a circularly symmetric shape and the nerve consisted of one fascicle. Extracellular recordings and stimulation did not take into account the electrode-electrolyte interface (see [25] for its effect on stimulation efficiency).

### 2.5. Axon Models

We used the original Hodgkin-Huxley parameters [26] for unmyelinated axons. Myelinated ones were based on the model of McIntyre et al.[27] that has originally been developed for peripheral motor fibers with thicker diameters (5.7 - 16.0 µm). To match the thinner axons found in the PNS (0.2 - 3 µm), we extrapolated all diameter dependent parameters to smaller diameters as shown in figure 3. Neither model is claimed to exactly match the properties of single neurons found in the PNS, but can rather be understood as a generic implementation, from which parameters to match specific datasets can be fine-tuned.

**Figure 3:**
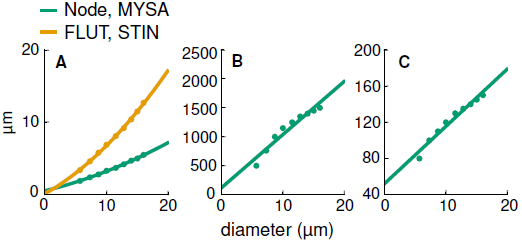
Linear and quadratic fits were used to extrapolate the parameters of myelinated axons to smaller diameters. (A) Diameters of all segments – nodes, MYSA (myelin attachment segments), and paranodal elements FLUT and STIN (see [27] for more information on the model) – were fit quadratically to prevent negative values. Node distance (B) and number of myelin sheaths (C) was extrapolated linearly.

### 2.6. Generation of Axonal Geometry

Axons in peripheral nerves are not perfectly straight, but instead follow the nerve path with a certain degree of tortuosity. To model this in our simulation – without defining the geometry for each fiber manually – we iteratively placed straight axon segments along a previously defined bundle guide, also composed of longer straight segments. In each step, the axon segment direction a_*i*_ was calculated as

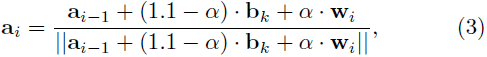

based on the corresponding bundle guide segment direction **b**_*k*_ (in general *k ≤ n* as bundle guide segments were longer than axon segments), the last axon segment direction a_*i*-1_ and a random component perpendicular to the bundle guide segment direction **w**_*i*_. All vectors have unit length. The parameter *α ∈* [0, 1] regulates the tortuosity of the axon and can, together with the distribution of ||**w**||, be fitted to geometries measured by microscopy. The factor (1.1 *-α*), rather than (1 *-α*), was chosen to maintain forward axon growth. See appendix A for the exact implementation of w_*i*_ to insure that axons stay within the nerve.

To fit our placing algorithm result to realistic axon trajectories, fibers in microscopy images were manually traced and segmented into straight sections of length 15 µm. For all traced axons of one nerve, the normed difference in direction between consecutive segments *c* = ||**a**_*i*_ − **a**_*i*+1_|| was calculated. We then compared the *c*-distribution of imaged, traced axons to the ones obtained from artificial fibres placed at different tortuosity coefficients α and ||**w**||-distributions to select the best fit. For details see appendix B.

### 2.7. Extracellular Potentials

Recordings from peripheral nerves capture changes in potential of the extracellular medium caused by membrane currents. To calculate those changes in `PNPy`, axon segments were interpreted as point current sources, each causing a potential change in the whole medium; see figure 4. Potentials of all current sources were superposed. Building on the the electro-quasistatic approximation of the Maxwell equations, combined with pure resistivity, time and space can be separated in the potential calculation:

**Figure 4:**
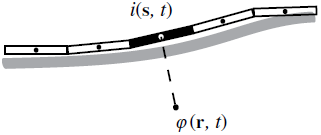
Axon segments can be interpreted as current point sources. The extracellular potential *ϕ* at position **r** caused by a current *i*(s, *t*) at position s is determined by current time course scaled with a static potential depending on the extracellular space and the spatial relation between source and receiver position.

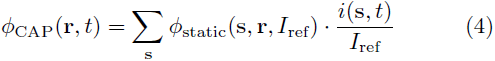

The static potentials caused by point current sources of reference current *I*_ref_ at source positions s, recorded at receiver position **r** were each scaled by the temporally varying axon segment membrane current *i*(*s,t*) and the resulting signals from all sources were added to obtain the extracellular potential time course _CAP_ at receiver position **r**.

Extracellular stimulation follows exactly the same principle, with stimulation electrodes modeled as assemblies of point current sources and axon segments as potential receivers.

### 2.8. Homogeneous Media

If the medium is assumed to be homogeneous with a constant conductivity *σ*, the potential *ϕ* (**r**, *t*) at **r** caused by a point source of current *i*(s, *t*) at s can be analytically written (see Malmivuo and Plonsey [28, Chapter 8] or [29] for reference) as

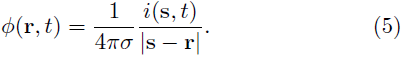

`PNPy` implements the homogeneous case as `PNPy.Extra cellular.homogeneous`.

### 2.9. Radially Inhomogeneous Media

As the medium surrounding the axons in peripheral nerves is anisotropic and inhomogeneous, the homogeneous assumption is not appropriate. Consequently, no exact analytical solution for the potential caused by a point current source exists and numerical methods become necessary. In order to reduce computational load, we took advantage of the separation of time and space in the potential calculation (4) and of existing symmetries in the extracellular space geometry. We precomputed potential fields using finite element method (FEM) software once for each unique point source position, then imported them into `PNPy`. This means that the computationally expensive field calculation only had to be carried out once per extracellular medium geometry. To insure the feasibility of this approach, the extracellular space was modeled using the simplified geometry shown in the upper drawing of figure 5, with conductivities set to the values given in table1. Conductivity only varied along the radial coordinate perpendicular to the bundle direction in this case, hence we refer to it as a radially inhomogeneous medium. Shifts of the source position along the nerve and rotations around its center are equivalent to shifting and turning the field. As a consequence, a very limited set of source positions, each for a distinct radius, completely characterises the field of any axon segment or stimulation electrode within the range of radii.

**Figure 5:**
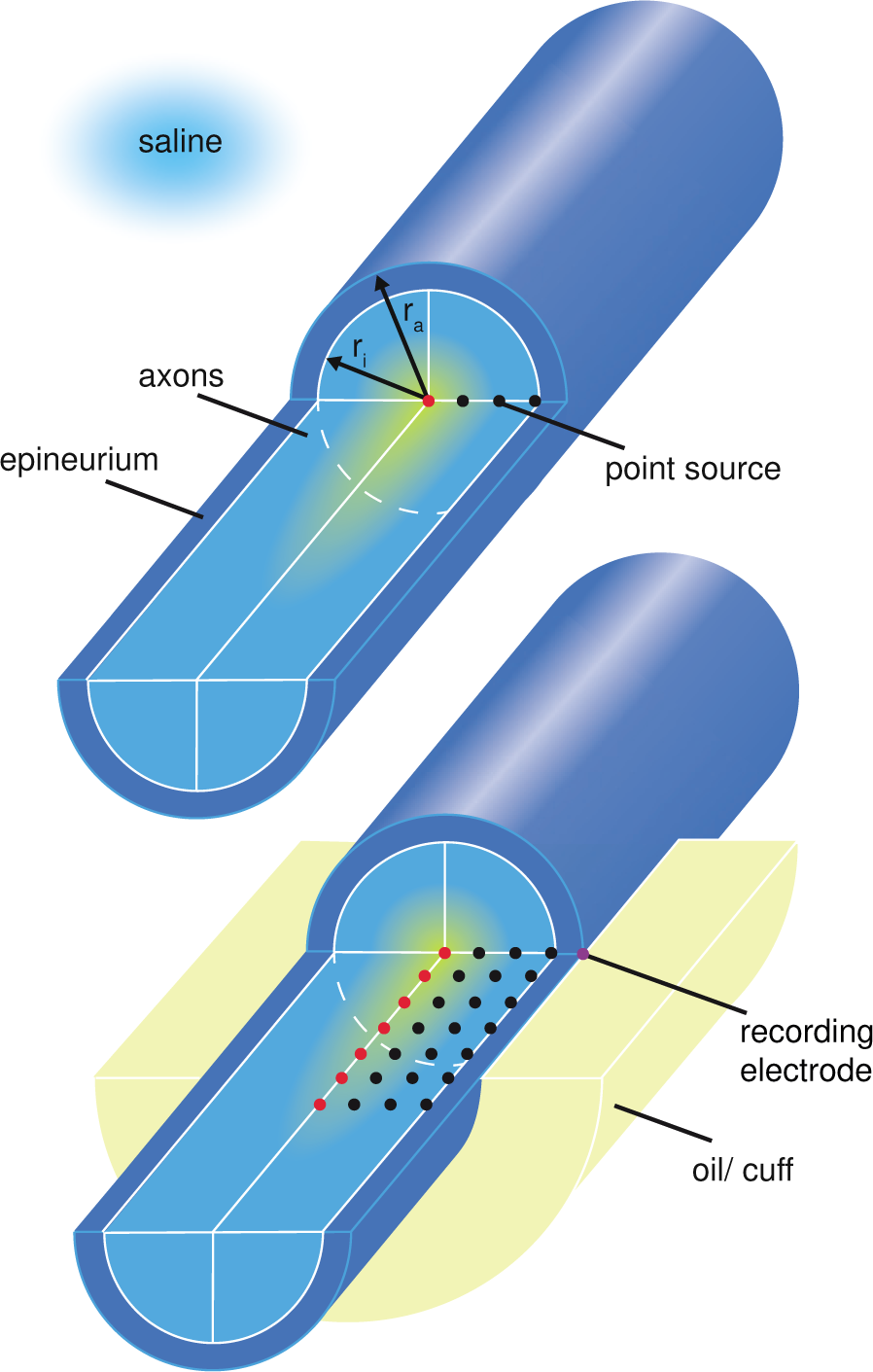
A circularly symmetric geometry makes import of precomputed potential fields feasible – if the medium is only radially inhomogeneous (upper drawing). A longitudinal inhomogeneity requires much more precomputed simulations (lower drawing). The nerve is modeled as axons (white matter) surrounded by the epineurium. The positions of exemplary current point sources, each generating one potential field, are shown. For radially inhomogeneous media, a line of sources does characterise all unique fields. For longitudinal inhomogeneities, potential fields for a two-dimensional array of point current sources need to be precomputed.

We calculated voltage fields *ϕ* (*x, y, z, r*) for different radial point source displacements *r* in the the FEM solver COMSOL Multiphysics 4.3. Due to our assumptions concerning the medium, steady state simulations were sufficient (separation of time and space). The static voltage fields were exported on a grid of *x ∈ -*1.5:0.015:1.5mm, *y ∈* 0:0.015:1.5mm, *z ∈* 0:0.03:30mm (start:step-size:end) where *z* is the longitudinal nerve axis and source positions are placed along *x*. The fields were imported in `PNPy` as a linear 4D spline interpolator. `PNPy` afterwards scales the static potentials with current time courses as given in (4) with *I*_ref_ set to 1 nA in COMSOL. The corresponding mechanism in `PNPy` is `PNPy.Extracellular.precomputedFEM`.

### 2.10. Longitudinally Inhomogeneous Media

In electrophysiological experiments, the nerve does not usually stay in its natural surrounding tissue. Instead, to strengthen the recorded or stimulated voltage, a cuff or a mineral oil bath increases the extracellular resistivity. The medium is in this case no longer longitudinally homogeneous, and any longitudinal shift in current source position will result in a different potential field. For stimulation, the current source (stimulation electrode) position can be fixed and the precomputation of very few potential fields, each for one electrode radius, characterises the effect of the electrode completely. For recordings, however, the longitudinal source position necessarily varies, as the axon segments extend through the nerve. Therefore, to cover all unique axon segment potential fields, a 2D-array of source positions distributed along both radial and longitudinal direction must be precomputed, as shown in figure 5. Note that without circular symmetry, a volume of source positions would need to be simulated, making the precomputation infeasible.

We found that for recording, a reasonable number of current source positions (∼20, each using about 40MB of memory) could not abolish interpolation errors between fields from longitudinally adjacent source positions, causing artifacts in the extracellular action potentials. To generate recordings without artifacts, a smooth analytic transfer function between point current source position and potential in the cuff was fit to FEM model results. Details are given in appendix C. This transfer function served in `PNPy` as a variant of `PNPy.Extracellular.analytic`.

## 3. Results

### 3.1. Axon models

For thin (< 1 µm) myelinated axons, extrapolated parameters from the McIntyre model [27] yielded bursting behavior. To prevent this, the potassium channel density at the nodes was increased by a factor of 1.5. Node size reduction with diameter achieved the same effect but is not observed [31, 32]. Potassium channels in the paranodal regions (not included in the original model) have been observed physiologically [33, 34] but their integration in the model could not abolish bursting. Myelinated conduction velocity (CV) fit experimental data well (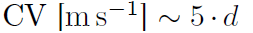 with diameter *d* in µm). Unmyelinated axons based on Hodgkin-Huxley channels had very low conduction velocities, 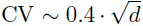, in comparison with expected values of around.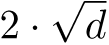 [35].

As membrane current directly shapes extracellular potential recordings, figure 6 compares the membrane current for both axon types over time before we cover single fibre action potentials in the next section. Importantly, unmyelinated axons emitted more current and the signal shapes differed considerably. The unmyelinated current time course was smooth, whereas the myelinated one was more complex with a sharp peak and a long lasting recovery. The latter axons contain different segments (node, myelin attachment segment (MYSA), paranodal main segments (FLUT)) which all contribute to the overall current output and thereby caused the more complex shape. See the model of McIntyre et al. [27] for more details on section types.

**Figure 6:**
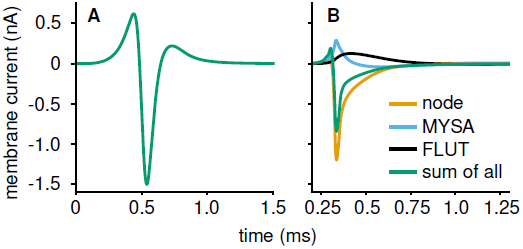
Unmyelinated axons (A) emit more current with a smoother time course than myelinated (B) ones. Both axons had a diameter of 3 µm. The corresponding segment lengths were 34.5 µm and 389 µm for unmyelinated and myelinated axon respectively, further increasing the difference in current per distance.

### 3.2. Profiles of Extracellular Media

The translation of membrane current to recorded potential by the extracellular medium is most strongly determined by the potential profile over longitudinal distance. To clarify its importance for the single fibre action potential, equation (6) explicitly states the SFAP calculation as a sum of single segment contributions for a straight axon running longitudinally along the nerve, here on the *z*-axis. If one assumes, as an extreme example, the Kronecker delta as a profile (*ϕ*(*z*) = *δ* (*z*)), the SFAP would have exactly the same time course as the membrane current. On the other hand a constant profile *ϕ* (*z*) = *c* will make the resulting SFAP vanish because of charge conservation (∫ *i*(*t*)*dt* = 0).

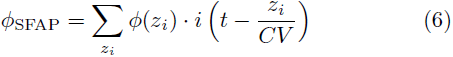

Further considering the shape of membrane currents as shown in figure 6, *ϕ*_SFAP_ in (6) will be maximal if positive and negative peaks of current add up constructively. To quantify when this happens, an active length *l*_a_ of an axon can be defined as

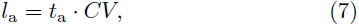

with *t*_a_ denoting the time during which an axon segment emits current of constant sign and *CV* the conduction velocity. Membrane current will be of the same sign over length *l*_a_. The fit between this length and the range of the profile 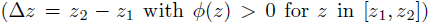 will determine the amplitude of the SFAP – in addition to a scaling factor depending on the absolute values of *ϕ* (*z*) in(6).

Building on those considerations, the profiles of our media can be compared. In figure 7, the strong impact of the cuff insulation is obvious. The potential profile became smooth, stretched out in space. The thin nerve surrounded by an insulation acted as two parallel resistors, causing a linear characteristic. For radial displacements of the source towards the electrode, a sharp peak emerged (see also figure C.1). We expect fast conducting axons with long *l*_a_ to best match this large range profile. The other two media had a different, much narrower characteristic. Radial inhomogeneities produced a slightly smoother potential profile compared to the homogeneous medium but differences remained small. Both profiles decayed a lot steeper with longitudinal distance than in the cuff and were therefore expected to better suit slower conducting axons with a shorter *l*_a_.

**Figure 7:**
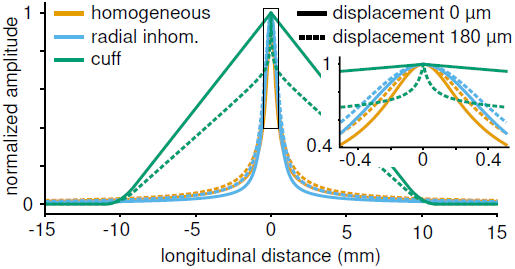
Compared to the homogeneous and radially inhomogeneous extracellular media the cuff insulation caused a much softer and strikingly linear characteristic.

### 3.3. Extracellular Single Fiber Action Potentials

We compared the effect of different extracellular media on the resulting SFAPs, results are shown in figure 8. Recordings from unmyelinated fibres mainly differed in amplitude and slightly in shape across media. Insulating the nerve with a cuff increased the potential by a factor of about ten and caused a narrower signal shape. In addition, an entrance and an exit peak at the sides of the cuff arose. The radially inhomogeneous medium slightly stretched the action potential in time which can be explained by the preference of current to flow along the nerve rather than transversally (compare to profile in figure 7).

**Figure 8:**
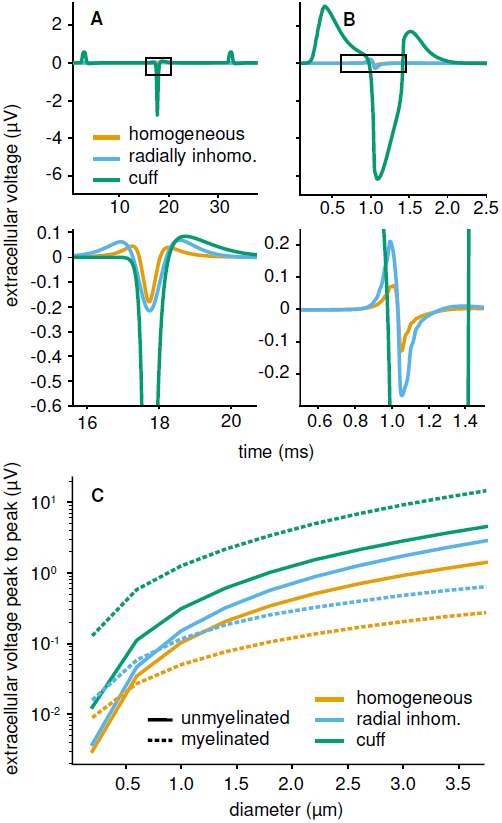
Unmyelinated and myelinated SFAPs showed different sensitivities towards the extracellular space. (A) The main peak of unmyelinated fibers mostly varied in amplitude over media, not in shape. In cuff insulated nerves, additional side peaks emerged.(B) Myelinated fibres produced much higher and longer lasting SFAPs in the cuff insulated medium. Both axons had diameter 3 µm, were placed centrally within the nerve and recorded by a circular monopolar electrode with radius 235 µm. Conductivity of the homogeneous medium was set to 1 S m^*-*^1. Lower row shows zoomed-in plots. (C) The amplitude boost achieved by cuff insulation was stronger for myelinated than for unmyelinated axons over the whole diameter range. For the other two media, unmyelinated SFAPs produced stronger SFAP amplitudes at diameters above 0.5 and 1 µm respectively.

The SFAP of myelinated axons was even more strongly affected when insulating the nerve. While the difference between homogeneous and radially inhomogeneous medium remained small, SFAP amplitude increased by a factor of about 20 in the cuff and shape was changed radically. The recorded signal lasted longer and had (as for unmyelinated axons) a negative main and two positive entrance and exit peaks.

The cuff insulation changed SFAP amplitude out of two reasons. One is the increased extracellular resistance. Current can not freely dissipate into the surrounding tissue but needs to flow along the thin nerve. As membrane current was modeled to be independent of the medium, an increase in extracellular resistance equaled an increase in extracellular potential. The second reason – that can explain the difference in amplitude gain between fiber types – is the match of active length (as defined in (7)) and cuff dimension (equal to range of the profile; 20 mm in this case) as detailed in section 3.2. For a myelinated axon of diameter 3 µm the active length evaluated to approximately 0.5 ms *·* 15 m s^*-*^1= 7.5 mm, an unmyelinated axon of this diameter only had an active length of about 0.5 ms *·* 1 m s^*-*^1 = 0.5 mm. Figure 8C demonstrates the matching effect between myelinated axons and the cuff over all diameters. While the SFAP amplitude of unmyelinated axons was similar and even higher in homogeneous and radially inhomogeneous media (narrow profiles), myelinated fibres achieved much stronger amplitudes following cuff insulation (wide profile) – even though their membrane current output is substantially lower compared to unmyelinated axons (see figure 6).

### 3.4. Compound Action Potentials

For validation, we aimed at reproducing experimental recordings from the stimulated rat vagus nerve in `PNPy`. To this end we obtained diameter distributions and fibre counts from microscopy images [36] - see table 2 - and set the geometry of the nerve and the recording electrodes so as to match the experimental set-up. Outer and inner radius were set to 240 µm and 190 µm respectively; a circular bipolar electrode of radius 235 µm and pole distance 3 mm (20 points per pole) surrounded the nerve. Axons were placed centrally and were activated intracellularly; due to the difference in stimulation threshold between fibers types, the entire population of myelinated and only a small fraction (∼20% of 10,000) of unmyelinated axons was triggered. As unmyelinated fibers based on Hodgkin-Huxley channels had very low conduction velocities, we corrected their SFAP timings.

**Table 1:**
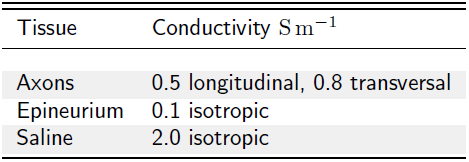
Conductivity of different tissues contained in a peripheral nerve [12, 30]

| Tissue | Conductivity $\text{S m}^{-1}$ |
| --- | --- |
| Axons | 0.5 longitudinal, 0.8 transversal |
| Epineurium | 0.1 isotropic |
| Saline | 2.0 isotropic |

**Table 2:** Axon number and properties set in the simulation for comparing model results with experimental recordings

| Type | # | diameter ($\mu\text{m}$) |
| --- | --- | --- |
| Unmyelinated | 2000 | $\in (0.2, 1.52)$ distribution from [36] |
| Myelinated | 200 | $\sim \mathcal{N}(2.3, 0.5)$ [36] |

Figure 9 demonstrates a reasonable agreement between simulation and experiment in the time domain. This only held for the cuff insulated medium – homogeneous and radially inhomogeneous media led to very low extracellular potential amplitudes as already observed for SFAPs and predicted from their lower tissue resistance.

**Figure 9:**
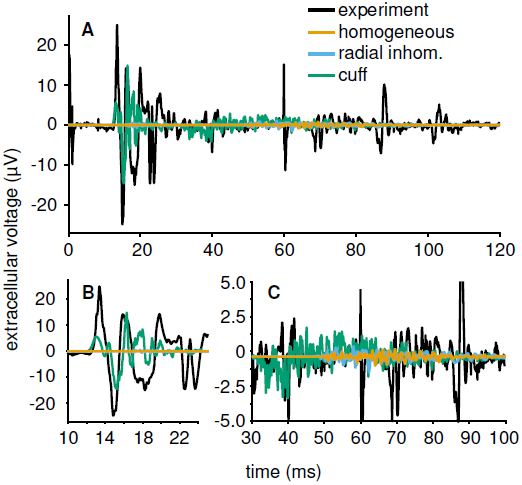
(A) The simulated compound action potential in the cuff medium approaches the experimental recording well. As expected, homogeneous and radially inhomogeneous media lead to much weaker signal amplitudes. See table 2 for axon properties. Distance between stimulation site and bipolar electrode (3 mm pole distance, 235 µm radius) was 8 cm with a length variability of 10%.All axons were activated by intracellular stimulation. The timing of unmyelinated SFAPs was adapted to regular conduction velocity values assumed in mammalian peripheral nerves (CV = 1.4 *√d*, CV in m s^-1^, d in µm). (B) The signal from myelinated fibers, which arrive first, appears very similar to the experiment. (C) The unmyelinated signal segment is shorter than in the experiment but matches the amplitude except for single large peaks.

Especially the signal portion caused by myelinated fibres (figure 9B) matches the experiment well in peak amplitudes, number of zero crossings and overall duration. Unmyelinated fibres (figure 9C) also produced a CAP comparable to the experiment in both amplitude and timing. However, differences existed: the simulated stream of action potentials ceased earlier and did not reproduce some strong late peaks observed in the experiment. Such late signal components could be explained either by lower conduction velocities or by longer fibres. In order for axon length alone to account for this delay, some fibres needed to be at least 30%*§* longer than the nerve - which is highly unlikely. The more probable explanation for such late high amplitude peaks would be an axon group of low conduction velocity and very similar fibre length whose single fibre contributions superpose constructively at the electrode; possibly amplified by an eccentric positioning within the nerve.

In the frequency domain (figure 10), the difference in fit to data between myelinated and unmyelinated fibers was confirmed. The spectrum of the unmyelinated signal proportion in our experimental data (black lines in figure 10A) had an overall flat profile with a main peak (lower plot) at around 500 Hz. This characteristic was approached to a certain extent by our model. The simulated spectrum of the cuff insulated nerve had a later peak at around 1 kHz but still followed the characteristic of the experiment well between 0 to 6 kHz before decaying further below -20 dB from there. We surmise that the high frequency content of the experimental data may be be caused by high frequency noise from the recording process. Meaningful, spike-event related signal components from experimental recordings usually stay below 2 kHz [20].

**Figure 10:**
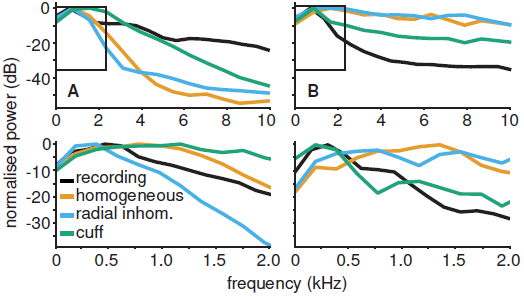
In the frequency domain, simulation and experiment did not match equally well for both fibre types. (A) For unmyelinated axons, the simulation did not perfectly approach the experimental spectrum in any medium with best results for the cuff. (B) The simulated frequency characteristic of myelinated axons in the cuff insulated medium was close to reality.

The experimental spectrum of myelinated fibers (figure 10B) was dominated by low frequency power below 2 kHz with an early peak at about 300 Hz. Our simulation result clearly matched this characteristic for frequencies below 2 kHz. Again only the cuff insulated medium model achieved this fit. The other two media led to a flat characteristic with a large amount of high frequency power.

In conclusion the experimentally obtained frequency characteristic of unmyelinated axons was reasonably, but not perfectly matched by our simulation. For myelinated fibres, we approached the experimental spectrum well as expected from the match in the time domain.

### 3.5. Fitting Axon Tortuosity to Experimental Data

In order to obtain axon shapes close to reality, we compared the distributions of axon segment direction changes *c* as detailed in methods section 2.2 for imaged mouse sciatic and vagus nerve – see figure 11 – and for our simulation at different parameters (||**w**||-distribution and *α*). A Gaussian ||**w**||-distribution produced *c*-distributions most similar to microscopy data (figure 12A). The sciatic nerve then corresponded to an *α*-value of about 0.6, the vagus nerve had a wider *c*-distribution as its axons were curvier, corresponding to a higher *α*.

**Figure 11:**
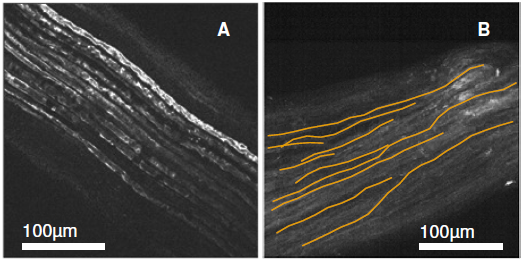
Fluorescence microscopy images of the mouse sciatic and vagus nerve both show slight tortuosity in their axon trajectories. (A) The thick myelinated fibers in the sciatic nerve appear very parallel. (B) The thinner axons in the vagus take a more curvy trajectory. Several manually traced fibers used to fit the model are highlighted in orange.

**Figure 12:**
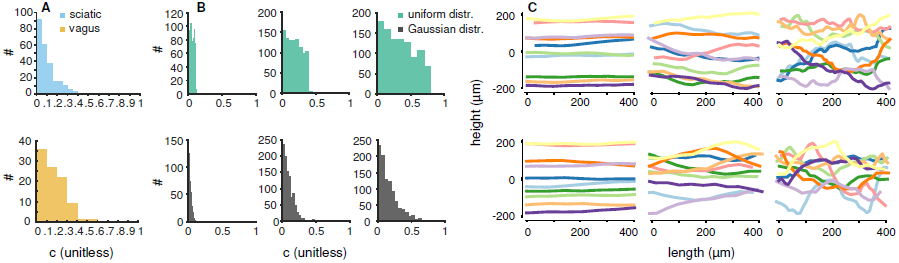
The axon placing algorithm was fit to tortuosity of microscopy imaged fibres. (A) The higher tortuosity observed in the vagus nerve is visible from the wider distribution of segment direction changes (*c*-values) compared to the sciatic nerve. (B) A normally distributed amplitude of the random component ||**w**|| in (3) better reflects the microscopy measured *c*-distributions in(A). Plots for s of 0.2, 0.6, and 1.0, respectively. Uniform ||**w**||-distributions result in a slightly skewed *c*-distribution because of the translation between ||**w**|| and *c* given in (B.1). Still, higher values are overrepresented. The normal distribution captures the dominance of low direction changes combined with rare high *c*-values. (C) When comparing the trajectories from both uniform (upper plot) and Gaussian (lower plot) at the same -values of 0.2, 0.6, and 1.0, it can be seen that the normal distribution of random vector length ||**w**|| leads to both a slightly smoother trajectory and rare strong direction changes, especially for high *α*-values.

### 3.6. Recording from Tortuous Axons

A more complex axon trajectory caused more complex SFAPs, as can be seen in figure 13. Unmyelinated SFAPs were especially sensitive to tortuosity. They developed complex, long lasting signals, especially in homogeneous and radially inhomogeneous media. When insulating the nerve, the amplitude of the main SFAP peak became very weak at high tortuosity while many small side peaks arose, giving the signal a noisy appearance. Myelinated fibers were more robust to tortuosity – their SFAP shape remained invariant at low and medium *α*-values. Only high degrees of tortuosity could change signal timing and shape; as for unmyelinated axons, the cuff isolated medium let the signal become noisy.

**Figure 13:**
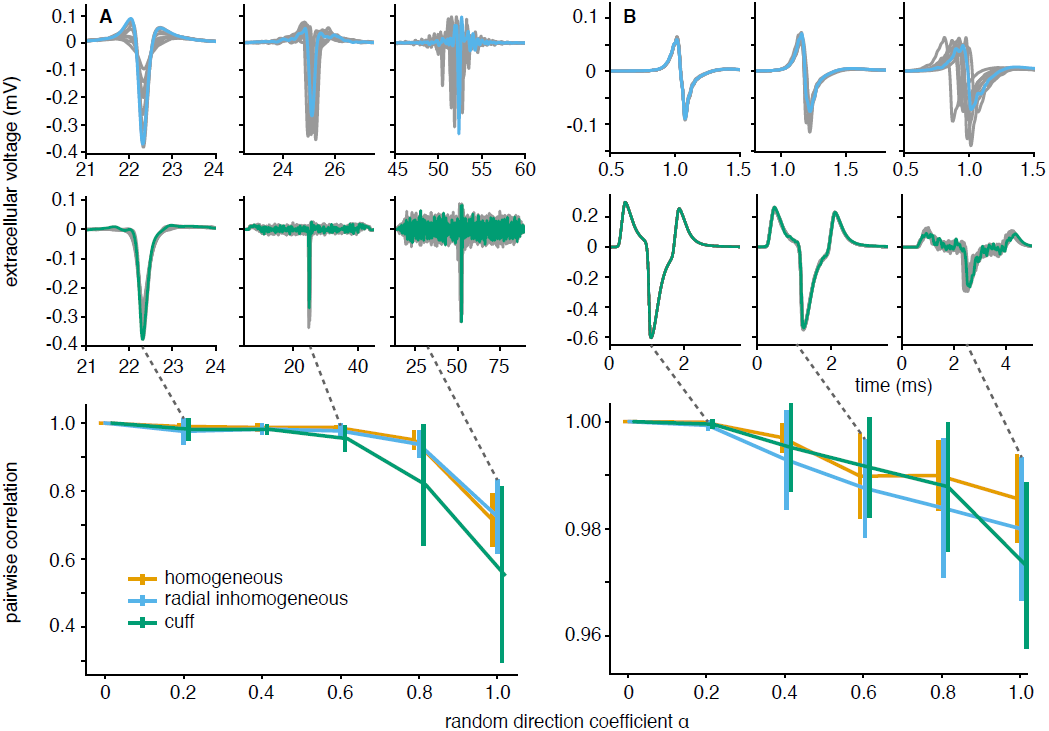
Unmyelinated axons were more sensitive to tortuousity in their SFAP shapes than myelinated ones. Tortuosity parameter α was set to 0.2, 0.6, and 1.0 for the signals shown in the first two rows. Gray lines correspond to SFAPs of different trials (axon geometries) at the same parameters. (A) Unmyelinated axons produced SFAPs differing both in timing and shape for the non-insulated nerve (radially inhomogeneous medium), even for small *α*-values of 0.2. In the cuff-insulated nerve (middle row) their signals became noisy at low *α*-values and the main SFAP peak almost disappeared for α = 1.0. (B) Myelinated axons mostly differed in timing in the radially inhomogeneous extracellular space, and not as much in shape. In the cuff, noisiness only arose at high tortuosity values. In the lower plots, the mean maximum pairwise cross-correlation gives a quantitative confirmation of the higher susceptibility of unmyelinated axons to change their SFAP shape in the presence of tortuosity. Note the different ordinate scales.

The overall effect of tortuosity to change SFAP shape can be understood by looking at equation (6) and changing it as in (9) where *s* is the distance along the axon. The longitudinal distance *z*(*s*) along the nerve becomes a function of *s*, shaped by tortuosity. Differences in the potential *ϕ* depending on the radial displacement of the axon were neglected here. The potential profiles of the extracellular media (see figure 7) are then both stretched (*z*(*s*) ≤ *s*) and distorted in a degree dependent on tortuosity. Different axons show different susceptibilities to this distortion because of their different active lengths. If the active length is large compared to the spatial frequency of the tortuosity-induced profile distortion, variabilities in *ϕ*(*z*(*s*)) are shadowed. Axons with shorter active length respond to those variabilities making their SFAPs noisier. This explains the difference in susceptibility between axon types.

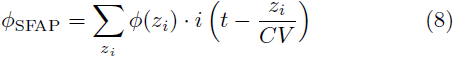

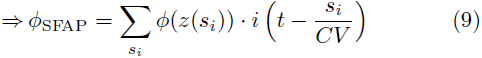

To quantify the influence of α on the heterogeneity of SFAP shape, we calculated the pairwise cross – correlation

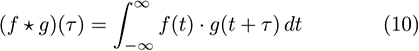

between normalized SFAP waveforms *s*_*α;i*_ from repeated simulation runs while keeping α, fibre type, and medium unchanged. The mean maximum cross-correlation over all waveform pairs described shape homogeneity:

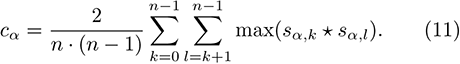

Figure 13 confirms that a higher *α* caused higher differences in shape (lower *c*_*α*_). As expected from the time course, myelinated SFAPs remained similar even for large s while unmyelinated ones lost their similarity. Note that this measure does take into account differences in timing or amplitude.

### 3.7. Stimulation of Tortuous Axons

Not only the recording from but also the stimulation of axons is influenced by their trajectory. Figure 14 shows that firstly, regardless of tortuosity, unmyelinated axons had much higher stimulation thresholds than did myelinated ones. Second, unmyelinated fibres had an optimal stimulation current with a smooth decrease in stimulation efficiency for higher and lower current amplitudes.

**Figure 14:**
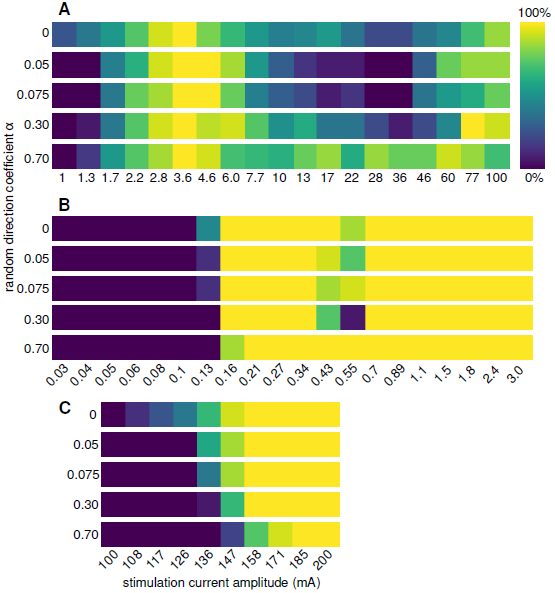
Unmyelinated axons have higher stimulation thresholds and are activated less reliably than myelinated ones. Both bundles consisted of 15 axons with diameter 3 µm and were stimulated with a bipolar electrode of radius 235 µm and pole distance 1 mm using a biphasic pulse of frequency 1 kHz, duration 1 ms and duty cycle 0.5. The extracellular medium was a nerve of diameter 240 µm bathed in oil. (A) Unmyelinated axons started to be activated at 1 mA and showed a peak in activation ratio at about 3 mA. (B) Myelinated fibers had a sharp activation threshold at a much lower current of about 0.15 mA and stayed activated for higher currents. Only when incrementing the stimulation current in very small steps of about 10 nA (C) a slight tortuosity-induced increase in stimulation threshold became visible for them as well.

Tortuosity affected the activation ratio of unmyelinated axons stronger than it did for myelinated ones. In the low amplitude range (< 3 mA), perfectly straight axons are activated best. In higher current regimes, very tortuous unmyelinated axons were the most consistently triggered. Stimulation of myelinated axons on the other hand was successful from low amplitudes of about 150 nA and at almost any higher current at all degrees of tortuosity. In figure 14C a minor increase in stimulation threshold with tortuosity becomes visible.

## 4. Discussion

The model that we have proposed here for the first time integrates compartmental axon models and numerically solved extracellular space models into a single simulation framework. To make the import of precomputed voltage fields feasible, the modeled media needed to fulfill certain constraints. One was the geometry that had to be circularly symmetric. Another constraint concerned material properties. Displacement currents and therefore frequency dependence of the tissues was not accounted for. Such frequency dependence certainly exists to a certain extent. It can arise from macroscopic structures at constant material properties (dielectric constant *ϵ* and conductivity *σ*) – the epineurium can for instance act as a capacitor. In addition, polarisation at different microscopic levels [37,38] can render the material properties *ϵ* and *σ* themselves frequency dependent. Such dielectric dispersion is observed in most biological tissues [39].

In terms of axon geometry, we implemented a simple iterative placement mechanism that was fit to microscopy data. It enabled us to investigate the influence of tortuosity on recordings and on stimulation efficiency indicating that perfectly straight axons are an oversimplification. To our knowledge this is the first implementation of such automated shape generation for peripheral nerve models. Our simulation predicted that SFAPs become more complex with increasing tortuosity – an effect that is exploited by spike sorting algorithms that differentiate single units from their SFAP shape. Axons were positioned independently from another and as a next step, fibre trajectories could be correlated as observed in microscopy images. Such correlation might explain some of the stronger peaks in the experimental CAP that we could not reproduce. SFAPs from fibres with very similar trajectory and therefore conduction delays could superpose to high amplitudes.

The modular nature of our model allows for an easy comparison of different extracellular media. The long temporal extent of SFAPs in cuff-insulated nerves – especially for myelinated axons – makes differentiation of single fiber contributions difficult as overlaps are probable. Unmyelinated axons (and slow myelinated axons) had separate entrance and exit peaks for the cuff, also making the temporal localization of their SFAP less easy. Overall a cuff therefore increased amplitude but reduced recording precision. Further our simulation predicts that matching the range of the extracellular medium to the active length of the fiber could significantly improve signal quality.

One limitation of the current NEURON simulation is the unmyelinated axon model. Its conduction velocity was too low compared to that reported for mammalian axons. For the overall CAP, the velocity needed to be corrected. Still, the Hodgkin-Huxley parameters are the accepted standard model for unmyelinated axons and more detailed C-fibre models (e.g. [40]) do not achieve significantly higher conduction velocities either. Parameters of the current model such as membrane capacitance or intracellular resistivity could be adapted to reach the expected conduction velocity but we chose to leave them at their physiological values. Interestingly, late peaks observed in the experimental recording hint on the existence of very slowly conducting axons.

Several steps to improve the model beyond the mentioned limitations are imaginable. First, axons are currently simulated sequentially. For the simulation of closed loop systems interacting with peripheral nerves, the simultaneous simulation of all nerves would be preferable. Second, axon membrane sections only need to be simulated if they are either stimulated or recorded from extracellularly, otherwise the calculation of their highly uniform membrane processes is unnecessary and time consuming. In order to eliminate computational overhead, one could introduce an abstract layer into the simulation in which the position change of spikes along axons is computed based on a known conduction velocity profile. Only for axon segments relevant to stimulation or recording, would the full membrane process be simulated.

In conclusion, a unified computer model of a peripheral nerve was developed. It combined an efficient calculation of extracellular potentials in inhomogeneous media from precomputed potential fields with compartmental axon models in a convenient Python module. The model was validated against experimental data and used to investigate the effects of conductivity inhomogeneities on amplitude and frequency content as well as the influence of axon tortuosity on both recording and stimulation. We hope that the simulation framework presented here, `PNPy`, becomes a useful tool for researchers working on peripheral nerves, nerve stimulation, and its medical applications, and envision that the toolbox could be augmented by multiple branches, organ models, and a variety of specific axon models matched to fiber types found in different parts of the peripheral nervous system, to facilitate this.

## Acknowledgements

This work was funded by EPSRC grant EP/L016737/1 and Galvani Bioelectronics; we further acknowledge support from grant EP/N014529/1. We would also like to thank Peter Quicke, Subhojit Chakraborty, and June Kyu Hwang for the two-photon microscopy images and Thomas Knöpfel for the ChAT-Cre FLEX-VSFP 2.3 mice used to obtain them.

## Appendices

### A. Calculation of the Random Component of the Axon Placing Algorithm

The random vector **w**_*i*_ in (3) is split into an inward pointing radial **w**_rad_ and a tangential component **w**_tan_ (A.1), both weighted independently with a weight drawn from a distribution *𝒫* (A.2, A.3). *𝒫* can be either a uniform distribution *𝒰*(−1, 1) between −1 and 1 or a normal distribution *𝒩* (*µ, σ*) with *µ* = 0 and *σ* = 0.33 (sigma chosen to have 99.7% of all values in the range [−1, 1]). When the radial distance between axon segment and bundle guide *d* approaches the bundle radius *r*_bundle_, the radial component **w**_rad_ becomes more inward directed (A.3) and thereby ensures that the axon stays inside the nerve. One linear implementation of the correction factor *e* is shown in (A.4). The parameter *r*_corr_ defined the relative radius from which on the correction should begin, set to 0.7 in our simulation; *e*_max_, set to 2 by default in `PNPy`, limits the correction.

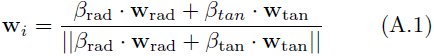

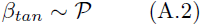

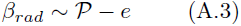

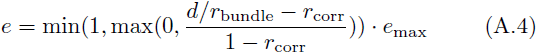

### B. Generation of Simulated *c* -Distributions

To directly translate ||**w**||-distributions (*𝒫*) to distributions of the normed difference in direction of consecutive axon segments *c* = ||**a**_*i*_ − **a**_*i*+1_|| projected onto a 2D-plane, we made the simplifying assumption that **b**_*k*_ and a_*i*_ are aligned. By doing so || **a**_*i*_ +(1.1 − *α*)*·***b**_*k*_||) (see equation (3)) becomes (2.1 *- α*) *· ||*a_*i*_||. Following figure B.1, it is easily shown that then ||**w**|| relates to *c* as

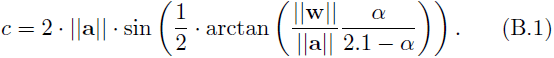

**Figure B.1:**
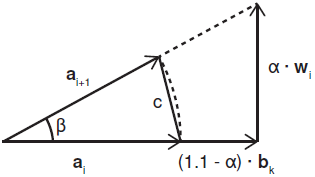
If bundle **b**_*k*_ and current axon segment a_*i*_ have a fixed relation, e.g. parallel, the expected distribution of segment direction differences *c* = ||**a**_*i*_ − **a**_*i*+1_|| can be easily obtained from the distribution of ||**w**|| (*𝒫*) by their geometrical relation.

### C. Fitted Cuff Transmission Function

For extracellular recording in a cuff, a transfer function between current point source position and the potential at an electrode longitudinally centrally placed in the cuff was fit. Input variables describe the spatial relation between source and receiver position. As apparent from figure C.1, the relation is strongly linear with an additional peak for low distances between current source and potential receiver – facilitating the fit of a transfer function.

The static potential was therefore described as a linear component *f*_lin_(*z*) plus a non-linear peak *f*_peak_(*z*). Equations (C.3) - (C.5) implement this characteristic for *ϕ* in mV with variables *r*_axon_ radial axon displacement in m, α angle between axon displacement direction and electrode perpendicular on the nerve center in rad and *z* longitudinal distance between electrode and axon in m. The transfer function is parametrized with *r*_1_ for the inner radius of the nerve in m, *a* and *b* for maximum peak amplitude and steepness, *c* for maximum of triangular component and *d* half the cuff width in m. The left and right borders of the *f*_lin_(*z*)-function were smoothed with a moving average of width *c/*20.

**Figure C.1:**
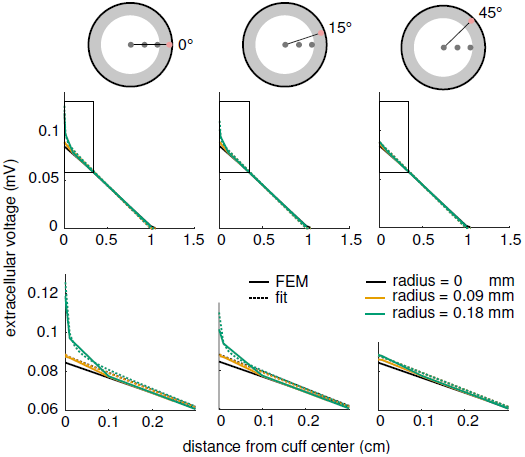
An analytic transmission function implements the relation between current source position and potential for recording in a cuff electrode. In the shown case, a nerve of diameter 480 µm in a cuff of 2 cm length was simulated in the FEM model. Functions are displayed for three different angles between the perpendiculars of source and electrode position onto the bundle guide respectively.

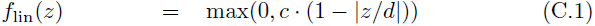

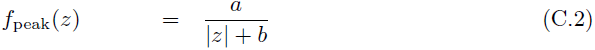

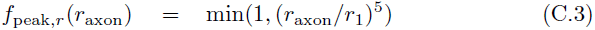

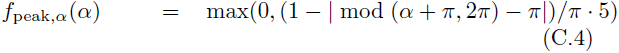

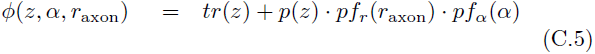

**Table C.1:** Parameters of the fitted transmission function for cuff recordings. Spatial input variables in m, angle in rad, output in mV.

| parameter | value |
| --- | --- |
| $r_1$ | $1.9 \cdot 10^{-4}$ |
| $a$ | $2.5 \cdot 10^{-9}$ |
| $b$ | $5 \cdot 10^{-5}$ |
| $c$ | $8.83 \cdot 10^{-4}$ |
| $d$ | 0.01 |

## Footnotes

‡ https://doi.org/10.5281/zenodo.579742

§ One late peak in the experiment occurs at 90 ms, the simulated signal decays around 70 ms. Simulated axon length needed to be increased by (90 ms - 70 ms)/70 ms *≈* 29% in order for the SFAP to be delayed accordingly.

